# Integration of Multiple Spatial Omics Modalities Reveals Unique Insights into Molecular Heterogeneity of Prostate Cancer

**DOI:** 10.1101/2023.08.28.555056

**Authors:** Wanqiu Zhang, Xander Spotbeen, Sebastiaan Vanuytven, Sam Kint, Tassiani Sarretto, Fabio Socciarelli, Katy Vandereyken, Jonas Dehairs, Jakub Idkowiak, David Wouters, Jose Ignacio Alvira Larizgoitia, Gabriele Partel, Alice Ly, Vincent de Laat, Maria José Q Mantas, Thomas Gevaert, Wout Devlies, Chui Yan Mah, Lisa M Butler, Massimo Loda, Steven Joniau, Bart De Moor, Alejandro Sifrim, Shane R. Ellis, Thierry Voet, Marc Claesen, Nico Verbeeck, Johannes V. Swinnen

**Affiliations:** STADIUS Center for Dynamical Systems, Signal Processing and Data Analytics, Department of Electrical Engineering (ESAT), KU Leuven, Kasteelpark Arenberg 10, 3001 Leuven, Belgium; Aspect Analytics NV, C-mine 12, 3600 Genk, Belgium; Laboratory of Lipid Metabolism and Cancer, KU Leuven and Leuven Cancer Institute (LKI), Leuven, Belgium; KU Leuven Institute for Single Cell Omics (LISCO), KU Leuven, Leuven, Belgium; Laboratory of Reproductive Genomics, Department of Human Genetics, KU Leuven, Leuven, Belgium; Laboratory of Stem Cells and Cancer, Université Libre de Bruxelles (ULB), Brussels, Belgium; Molecular Horizons and School of Chemistry and Molecular Bioscience, University of Wollongong, Northfields Ave, Wollongong, NSW 2522 Australia; Department of Pathology and Laboratory Medicine, Weill Cornell Medical College, New York, NY, USA; Department of Analytical Chemistry, Faculty of Chemical Technology, University of Pardubice, Pardubice, Czech Republic; Laboratory of Multi-omic Integrative Bioinformatics, Department of Human Genetics, KU Leuven, Leuven, Belgium; Department of Urology, University Hospitals Leuven, Leuven, Belgium; Department of Development and Regeneration, KU Leuven, Leuven, Belgium; South Australian Health and Medical Research Institute (SAHMRI), Adelaide, SA 5000, Australia; South Australian Immunogenomics Cancer Institute, University of Adelaide, Adelaide, SA 5005, Australia; Freemasons Centre for Male Health and Wellbeing, University of Adelaide, Adelaide, SA 5005, Australia

## Abstract

Recent advances in spatial omics methods are revolutionising biomedical research by enabling detailed molecular analyses of cells and their interactions in their native state. As most technologies capture only a specific type of molecules, there is an unmet need to enable integration of multiple spatial-omics datasets. This, however, presents several challenges as these analyses typically operate on separate tissue sections at disparate spatial resolutions. Here, we established a spatial multi-omics integration pipeline enabling co-registration and granularity matching, and applied it to integrate spatial transcriptomics, mass spectrometry-based lipidomics, single nucleus RNA-seq and histomorphological information from human prostate cancer patient samples. This approach revealed unique correlations between lipids and gene expression profiles that are linked to distinct cell populations and histopathological disease states and uncovered molecularly different subregions not discernible by morphology alone. By its ability to correlate datasets that span across the biomolecular and spatial scale, the application of this novel spatial multi-omics integration pipeline provides unprecedented insight into the intricate interplay between different classes of molecules in a tissue context. In addition, it has unique hypothesis-generating potential, and holds promise for applications in molecular pathology, biomarker and target discovery and other tissue-based research fields.

## Introduction

Recent developments in spatial-omics technologies have revolutionised tissue-based research, enabling a deeper understanding of the cellular architecture of tissues, cell-cell interactions as well as the intricate relationship between molecular identity and tissue structure. These technologies find broad applications in studying normal tissue development^1^, as well as in deciphering complex diseases like neurodegenerative disorders^2,3^ and different types of cancer^4–9^. To investigate the various classes of molecules (RNA, proteins, metabolites, lipids) in a spatial context, different technologies are applied, including *in situ* hybridization, *in situ* sequencing and *in situ* RNA capturing followed by *ex situ* sequencing for spatial transcriptomics^10–19^ (ST), histochemistry using fluorescently labelled antibodies^19–21^ or genetically encoded fluorescent protein tags^22,23^ for spatial proteomics, and mass spectrometry imaging (MSI) to study the distribution of peptides, metabolites, and lipids^24–29^.

The biological questions addressed by such platforms are often complex and involve molecular alterations at multiple levels of the gene expression cascade. As each of the spatial-omics technologies often captures only a partial aspect of a single molecular layer in a tissue, integration of spatial multi-omics (SMO) datasets that span across the biomolecular scale is essential to provide insight in the complex interplay between the different molecular layers and to better understand the intricacy of these biological processes, and their molecular drivers. Pipelines aiming to align and correlate these datasets encounter several obstacles, with the limited coverage of the molecular landscape being a prominent example. Moreover, distinct spatial technologies may not be compatible on a single tissue section, making the use of neighbouring tissue sections necessary^30^. As the structural organisation of tissues can differ between consecutive sections, and the experimental handling for the various platforms may result in tissue deformation, precise co-registration of adjacent datasets has proved difficult but is essential in order to align corresponding regions and prevent misinterpretation. This operation may be additionally complicated by the fact that many spatial technologies acquire data at different spatial resolutions, ranging from sub-cellular for microscopy-based technologies^31,32^ to 55 µm for the Visium ST platform^33^ and 1-100 µm for commercial MSI platforms^34^. Therefore, a granularity matching step is required to ensure that the spatial positioning of data points from the different modalities is reasonably equivalent. Finally, in spatial-omics studies, sensitivity, especially for detecting low-abundance molecules and achieving quantitative accuracy, poses further challenges. To address these limitations, spatial omics approaches can be complemented with bulk or single-cell omics technologies. Bulk-omics methods offer high coverage and sensitivity, enabling a more comprehensive and quantitative analysis of molecular profiles^35^, while single-cell omics provides insights into cellular heterogeneity within tissues, enhancing the resolution and sensitivity of spatial data^30,36^.

Recent studies have used a variety of different omics integration strategies, including co-registration and/or the use of a weighted nearest neighbour strategy to partially address the issues related to the combination of multiple spatial omics technologies^8,9,37,38^. Here, we have applied a novel spatial multi-omics pipeline (SMOx), which involves multi-point co-registration and Gaussian weighted granularity matching steps for more accurate data point matching, resulting in improved integration of various spatial omics modalities. Using this pipeline, we combined Visium ST (further referred to as ST) with MALDI-MSI coupled with post-ionisation (MALDI-2), which has increased sensitivity for detecting lipids compared to conventional MALDI^39,40^, and single-nucleus RNA sequencing (snRNA-seq) to perform cell-type deconvolution of ST data^41^ and high resolution pathology annotations. To explore the potential of this pipeline, we applied it to study human prostate cancer (PCa), which in view of its highly inter-and intra-tumoural heterogeneous nature^42^ is an ideal case study for the SMOx integration pipeline. In fact, PCa often exhibits multiple morphological disease states within a single tissue section, ranging from prostatic intraepithelial neoplasia (PIN) to various histological malignant states (ISUP grades) and cribriform, each representing distinct disease subtypes with unique biological characteristics and clinical outcomes^43^. Moreover, PCa development involves molecular changes across multiple omics classes. Besides alterations in genes and transcripts, prominent changes also occur in lipids, offering unique potential for novel therapeutic interventions^44,45^. Using PCa as a paradigm, we showcase the potential of our novel spatial multi-omics pipeline to uncover lipid-gene expression pairs that are linked to different cell populations and histopathological disease states not discernible by morphology alone and can aid in the annotation of histologically ambiguous regions. These findings illustrate the potential application of our data integration pipeline to diverse fields of tissue-based research, including molecular pathology and for the generation of novel research hypotheses.

## Results

### Establishment of a novel spatial multi-omics data integration pipeline (SMOx) and its application in prostate cancer

With the aim to more comprehensively map the molecular heterogeneity within tissues and to better understand the interplay between different classes of molecules in a tissue context, we established a spatial multi-omics pipeline (SMOx) and applied it to the field of PCa (**Fig. 1A)**. After collecting tumour and matching non-malignant biopsies from men that underwent a radical prostatectomy (n=8, **Supplementary Table 1**), 10 μm thick frozen tissue sections were cut. Neighbouring tissue sections were subjected to ST analysis using the Visium Spatial Gene Expression platform (n=16) and MALDI-2-MSI based lipidomics (n=16). From the ST analysis, we obtained curated expression data of 18,950 genes across 42,475 individual capture spots on the ST array slides (summarised across all samples). Only spots under the tissue sections were considered for downstream computational analysis. From the MSI data, peak picking identified 12,510 peaks across the samples. The average overall spectra of cancer and non-cancer samples are presented in **Supplementary** Fig. 1. High-resolution digital H&E images of the MSI sections were annotated by two independent expert uro-pathologists. Additionally, we collected three 100μm thick cryo-sections and isolated nuclei for snRNA-seq (n=13) from the remaining tissue directly adjacent to the cryo-sections used for ST and MSI analysis, to enable the identification and mapping of distinct cell populations across the tissue by applying the ‘cell2location’ deconvolution algorithm^46^.

**Figure 1:**
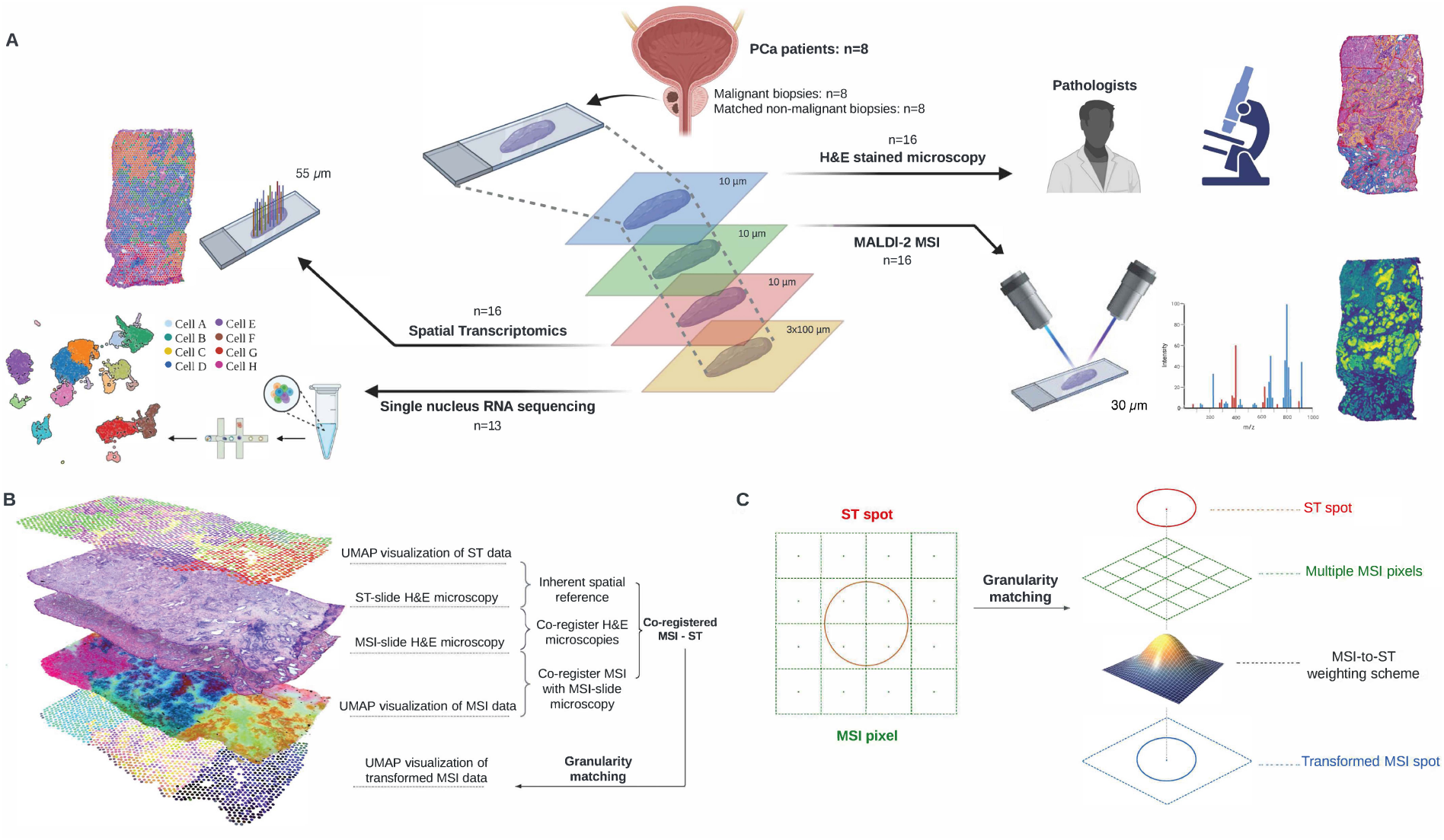
Study design and overview of the spatial multi-omics integration pipeline (SMOx). **A.** Schematic representation of the various datasets generated on prostate cancer samples and the applied integrated spatial multi-omics workflow. **B.** Strategy for co-registration and integration of different spatial multi-omics data layers generated on consecutive tissue sections. To link ST data with MSI data, UMAP visualizations of different modalities were co-registered in two steps based on the haematoxylin and eosin (H&E) stainings of the respective slides and MSI data, and a UMAP visualisation of transformed MSI data was generated after granularity matching. **C.** Overview of the granularity matching step to bridge the scale gap between MSI and ST data. Spatial readouts were aligned, and Gaussian weighting was applied to generate a transformed MSI spot.

To integrate the data from the ST and MSI spatial-omics modalities, which are generated on separate (neighbouring) tissue sections and are acquired at different spatial resolutions, the SMOx pipeline was designed to allow co-registration and granularity matching (**Fig. 1B-C**). As both ST and MSI routinely include H&E staining of tissue sections in their workflows^47–49^, we used the H&E-stained images to align the ST and MSI data. To facilitate cross-modality comparison and uncover inherent molecular patterns, the data from the different modalities were independently subjected to dimensionality reduction using uniform manifold approximation and projection (UMAP)^50^. The UMAP representation (i.e., mapped 3 dimensional embeddings) of the MSI data^51^ was used to guide the co-registration with the respective post-MSI H&E image. Subsequently, the MSI H&E image was co-registered to the H&E image of the ST Visium slide. Alignment of ST data with the respective H&E histological images was done automatically using the Space Ranger pipeline (10X Genomics), eliminating the need for manual co-registration^52^. This strategy of using H&E-stained images to align ST and MSI data reduces the complex multimodal registration to a mono-modal one between their respective H&E-stained images, to which each modality has previously been co-registered, and removes the selection effect of registering to a template image. Subsequently, a shared spatial coordinate system was created by linking the MSI data with the ST data. This approach produces a readout that can be considered a “virtual section”, between the sections used for ST and MSI analysis (**Fig. 1B**). Following co-registering of MSI and ST data, we addressed the disparity in spatial scales of data acquisition points through granularity matching. In this study, MSI was conducted at a spatial resolution (pixel size) of 30 µm, whereas the Visium assay has spot diameters of 55 µm with a 100 µm centre-to-centre distance between spots^33^. Consequently, each Visium spot corresponds to multiple MSI pixels with uneven amounts of coverage **(Fig. 1C)**. To match the granularity of the spatial readouts, we employed Gaussian weighting to the MSI pixels close to each ST spot after registration^53^. This procedure resulted in the generation of a comprehensive aggregated MSI feature spectrum linked to each ST spot, ensuring synchronised spatial readouts between MSI and ST data (**Fig. 1C**). The resulting co-registered and matched MSI-ST data structure can be interpreted as a data frame consisting of matched readouts for each Visium spot. Within this data frame, each row represents a single matched spot, while the columns encompass distinct dataset information at that specific spot, such as gene expression, lipid content, pathology annotations, deconvoluted cell types, etc. Thus, this integrated spatial multi-omics pipeline could link transcript-lipid pairs with tissue features.

### Application of the SMOx pipeline to identify correlated transcript-lipid pairs linked with histological annotations

Using our spatial multi-omics integration strategy, we explored the spatial correlation of transcripts and lipids in a PCa tissue section of a single patient (patient 929) as a proof of concept. The level of detail in the annotations created by the histopathologist on the digital H&E image, along with the transformation of annotations in the shared coordinate system, is depicted in Fig. 2A. In this tissue section, five histologically distinct regions could be discerned: benign epithelium, stroma, PIN, tumour ISUP 3 and immune cells. When UMAP embeddings are projected onto tissue locations, similar patterns emerge among the distinct spatial modalities, indicating that despite the diverse nature of spatial data readouts (genes, lipids, etc.) (Fig. 2B), they all contain relevant molecular information reflecting the distinct histopathological features in the sample.

**Figure 2.**
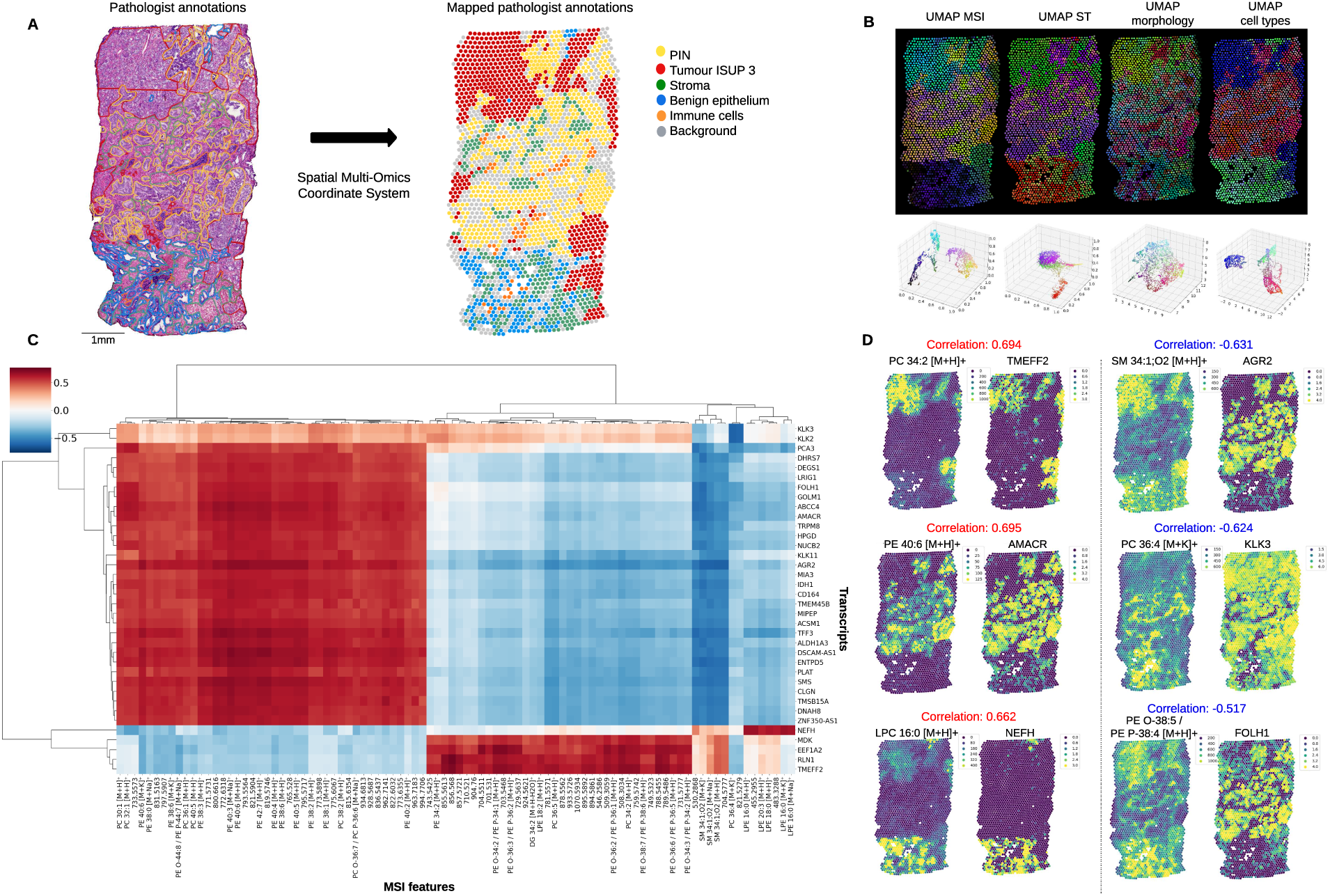
Integration of spatially resolved multi-omics reveals spatially correlated lipid-gene pairs in a prostate cancer sample. **A.** Highly detailed pathologist annotations of a representative prostate cancer sample (929_cancer) on the post-MSI H&E-stained tissue section and their spatial distribution projected on the shared coordinate system. **B.** UMAP representations of individual modalities including MS imaging data, ST data, morphological features extracted from pathology annotations, and cell types derived from snRNA-seq data after spot deconvolution. The top row represents a hyperspectral visualisation of the UMAP data on the shared coordinate system using the RGB colour scheme. The bottom row shows the scatter plots of the resulting 3-dimensional UMAP embeddings for each modality. **C.** Hierarchically clustered heatmap of the correlation matrix between MSI features and ST data. **D.** Spatial visualisation of representative positively and negatively correlated lipid-transcript pairs in the 929-cancer sample.

To link the different levels of molecular information, we conducted correlation analysis assessing the relationship between individual transcripts and MSI features (most of which are lipids), along with their spatial localization in the tissue (**Fig. 2C-D, Supplementary Fig. 2**). Among the positively correlated lipid-transcript pairs (correlation coefficient over 0.65), we observed PC 34:2 [M+H]^+^ and *TMEFF2* (a transmembrane proteoglycan with multiple roles in cancer^54^) as one of the most highly correlated pairs in regions annotated as tumour ISUP3. PIN included PE 40:6 [M+H]^+^ and *AMACR* (an established PCa marker^55^) as a highly correlated pair. LPE 16:0 [M+H]^+^ and *NEFH* (a neurofilament-encoding gene maintaining cell integrity) were identified in the benign epithelium region. In contrast, a notable spatial negative correlation (-0.631) was observed between the lipid SM 34:1;O2 [M+H]^+^ and the *AGR2* gene (a disulphide isomerase gene family implicated in tumour metastasis^56^), with their expression detected in tumour ISUP 3 and PIN regions, respectively. Interestingly, several of the identified transcripts listed in Fig. 2C encode enzymes that are directly involved in lipid metabolism, including DEGS1 (a sphingolipid desaturase^57^), HPGD (a 15-hydroxyprostaglandin dehydrogenase associated with PCa risk^58^), AMACR (an enzyme involved in peroxisomal beta-oxidation of branched-chain fatty acids and one of the most widely used histological markers upregulated in PCa^59^), and ACSM1 (a medium chain Acyl-CoA Synthetase that is overexpressed in PCa^60,61^). Importantly, these genes are up-regulated in areas annotated by pathologists as PIN and tumour ISUP3, in comparison to the regions annotated as benign epithelium. Many other transcripts are not directly implicated in lipid metabolism but are known to have a cell type-specific expression pattern and are differentially expressed upon cancer development. These include transcripts of genes *TMEFF2*, *KLK3* (encoding the prostate-specific antigen biomarker), *PCA3*, *AGR2*, and *NEFH*. These observations suggest that transcript-lipid correlations are associated with disease state and specific cell populations, and may reveal functional interactions.

### Identification of molecular expression patterns associated with histological annotations

To further explore the alignment of MSI data and ST expression patterns with pathology annotations, we applied non-negative matrix factorisation (NMF) to extract correlated features from the individual modalities of the integrated spatial multi-omics dataset of sample 929_cancer. The resulting correlation heatmap between MSI NMF components and pathology annotations demonstrated significant positive correlations for each pathological annotation (PIN, benign epithelium, tumour IUSP3 and stroma) except for immune cells (likely due to their greater cell type variability) (Fig. 3A **top panel**). The spatial distribution of these NMF components, along with pathology annotations within the common coordinate system are illustrated in Fig. 3B, accentuating the congruence between lipidome and histology. Similarly, significant positive correlations were observed between ST NMF components and the different annotated regions (including immune cells) (Fig. 3A **bottom panel** and Fig. 3B). For MSI NMF component 9, which highly correlates with PIN, the pseudo-spectrum of the biomolecular ion peaks that are involved in the expression of the MSI NMF component’s spatial patterns is shown, along with the gene expression profile of the PIN-associated ST NMF component 13 (**Fig 3A**). Integration of both spatial-omics modalities within the shared coordinate system and subsequent hierarchical clustering revealed strong correlations between specific NMF components of both modalities (Fig. 3C). Differentially expressed lipids and genes that contribute significantly to the molecular characteristics of the different annotated areas are listed in Fig. 3D. The high level of correlation provides further credence to a spatial relationship between readouts of the ST and MSI data, and overall agreement with the pathology annotations.

**Figure 3.**
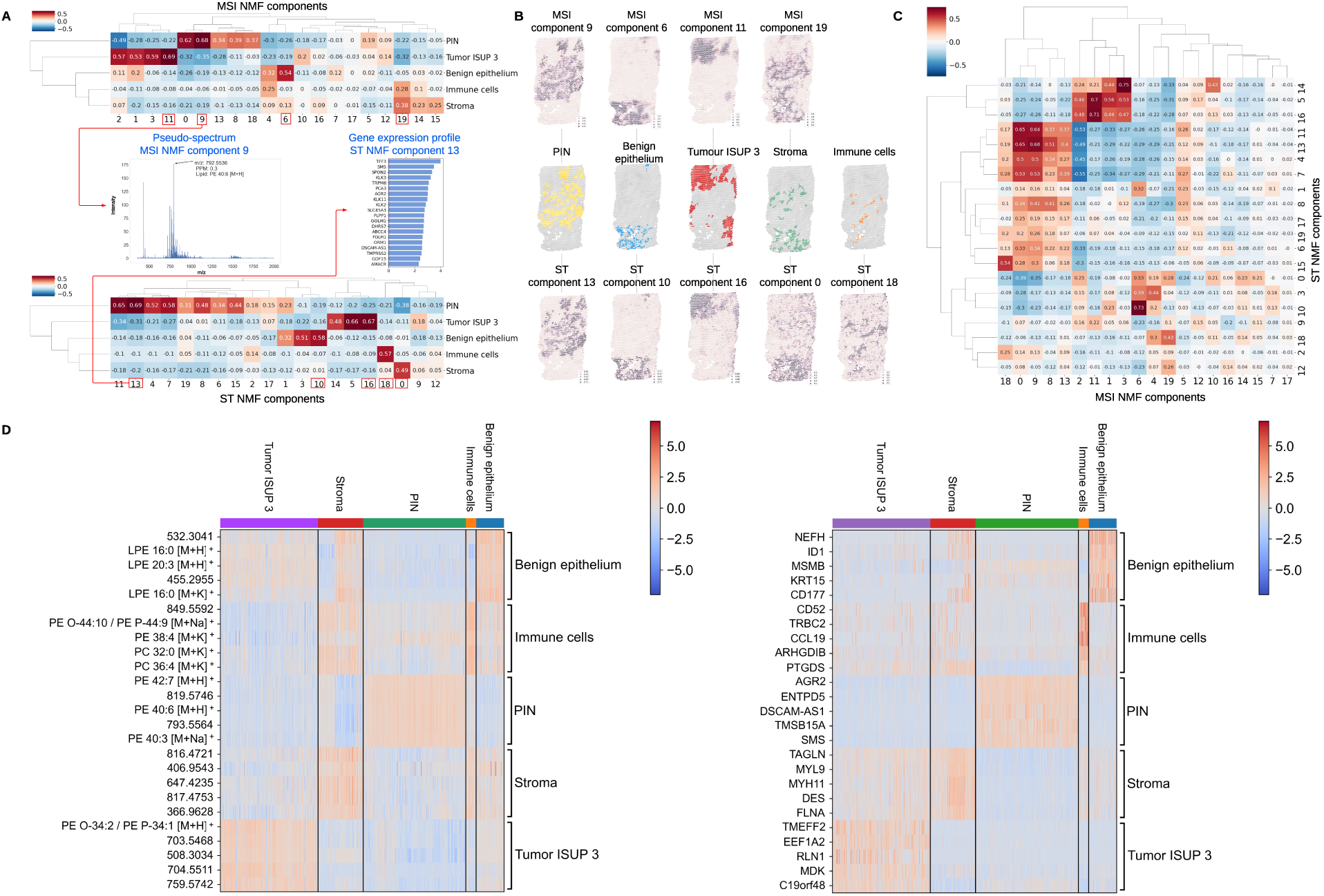
Correlation between histological and molecular profiling. **A.** Heatmap showing correlations between MSI NMF components and the five histologically distinct regions from sample 929_cancer. A representative pseudospectrum for MSI NMF component 9 and the most enriched genes of the correlating ST component 13 are shown. **B.** Spatial visualisation of representative MSI NMF components (top panels) correlating with ST NMF components (bottom panels) for each histological feature (middle panel). **C.** Hierarchically clustered heatmap of the overall correlations between MSI NMF and ST NMF components. Numerical values correspond to the respective correlation coefficient of each MSI NMF – ST NMF pair. **D.** Heatmaps illustrating MSI features (mostly lipids) (left panel) and genes (right panel) exhibiting the most significant differential expression within each of the five histological regions in the 929_cancer sample.

### Linking lipids and transcripts to specific cell types in the spatial coordinate system

Due to the granularity of the matched MSI-ST data set with a spatial resolution defined by the spatial-omics analysis techniques used (ST: 55 µm resolution; MSI 30 µm resolution), each spot in the spatial coordinate system corresponds to multiple cells, hampering the assignment of lipids and transcripts to specific cell types. To address this issue, we applied snRNA-seq-based spot deconvolution using the ‘cell2location’ deconvolution algorithm^46^, based on snRNA-seq data from a set of 100 *μ*m thick serial tissue sections neighbouring those utilised for ST. The resulting UMAP clusters were annotated based on curated marker genes^62–64^, leading to the identification of 14 distinct cell types (Fig. 4A). Of note, the TC1 (Tumour Cell 1) cell type was mainly enriched in both the tumour ISUP 3 and PIN region, whereas TC2 (Tumour Cell 2) was linked mainly to the PIN region only (Fig. 4B**-C**). Interestingly, cells with a club cell-specific gene signature, which have recently been associated with PCa carcinogenesis^63^, localised to sub-regions within the PIN-annotated areas. Correlation analysis between MSI and ST data from spots enriched in specific cell types revealed unique lipid profiles associated with a cell type along with marker genes and enzymes involved in lipid metabolism, as illustrated or TC1 and TC2 cells, club cells and macrophages (Fig. 4D). This analysis revealed that ISUP 3-associated TC1 cells and PIN-associated TC2 cells correlated with long chain polyunsaturated phospholipids (mainly PE), with an additional enrichment for hexosylceramides (HexCer) in the TC2 cell component. Cholesteryl esters (CE), mainly identified as [2M+H]+ ions^65^ and known to accumulate in lipid droplets, were associated with club cells. Macrophage marker genes, like *HLA-D* genes^66^, *LYZ*^67^ and *CXCL5*^68^, correlated best with ether lipids, CE and long polyunsaturated phosphatidylcholines (PC). Complementary gene signature analysis of lipid metabolism pathways revealed an overall high expression of genes linked to lipid metabolism pathways in luminal and TC2 cells, along with cell type-specific pathway activations (Fig. 4E). Examples include the upregulation of fatty acid biosynthesis and elongation in cancer cells (TC1 and/or TC2), steroid biosynthesis in luminal cells, and arachidonic acid metabolism in macrophages and mast cells, in line with expectations and data available in the literature^44,69,70^. Projection of these pathways in the UMAP representations confirmed these links and provided an additional layer of granularity, for instance revealing a specific subset of cells within the TC1 and TC2 clusters displaying elevated expression of genes involved in linoleic acid metabolism (Fig. 4F). These findings illustrate the benefit of linking lipids and transcripts to specific cell types in the spatial coordinate system.

**Figure 4.**
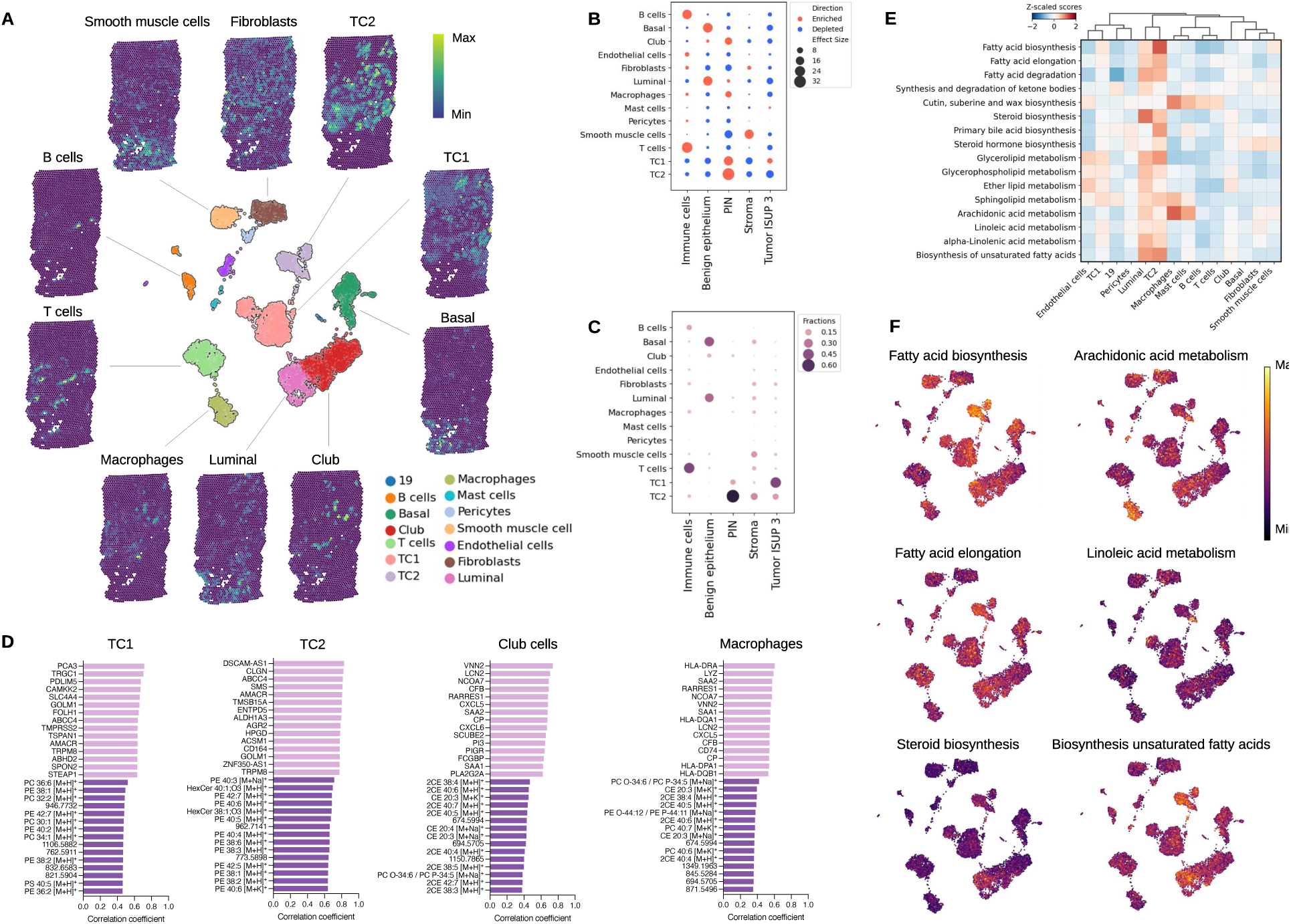
Spot deconvolution reveals cell type specific transcriptional and lipidomic profiles. **A.** UMAP representation of annotated snRNA-seq clusters and spatial localisation of the main cell types in the spatial transcriptomics data after spot deconvolution using cell2location^42^. TC1: tumour cluster 1, TC2: tumour cluster 2. **B.** Dot plot displaying the enrichment (orange) and depletion (blue) of cell types within the regions demarcated by the pathologist. **C.** Dot plot illustrating the relative abundance of a specific cell type within each pathologically annotated region. **D.** MSI features (pink) and transcripts (purple) with the highest positive correlations with TC1, TC2, club cells and macrophages. **E.** KEGG enrichment of differentially expressed genes involved in lipid metabolism per annotated cell type based on the snRNA-seq data. Colours indicate mean up-(red) or down-(blue) regulation of the pathway, while colour intensity indicates significance level. **F.** UMAP representation of the enrichment of different lipid metabolism pathways per cell type.

### Integrated data analysis reveals molecular heterogeneity in neoplastic disease areas

Whereas the spatial distributions of MSI NMF components and ST NMF components in sample 929_cancer overall matched well with the histological annotations, the region annotated as PIN by the pathologists, was captured by two NMF components in both modalities – MSI NMF component 9 and 18 and ST NMF components 13 and 15 (Fig. 5A**-B**). Additionally, snRNA-seq-based spot deconvolution revealed that one of the components of each modality (MSI NMF 9 and ST NMF 13) corresponded well with the spatial distribution of TC2 cells (Fig. 5C). The other component (MSI NMF component 18 and ST NMF component 15) aligned with a specific subregion of PIN corresponding to the spatial distribution of epithelial club cells and macrophages. Examination of the H&E stainings of these latter regions confirmed the presence of a neoplastic gland-like structure. These observations underscore the potential of the integrated data analysis approach to reveal molecular heterogeneity in neoplastic disease areas that might not be discernible through morphological examination alone.

**Figure 5.**
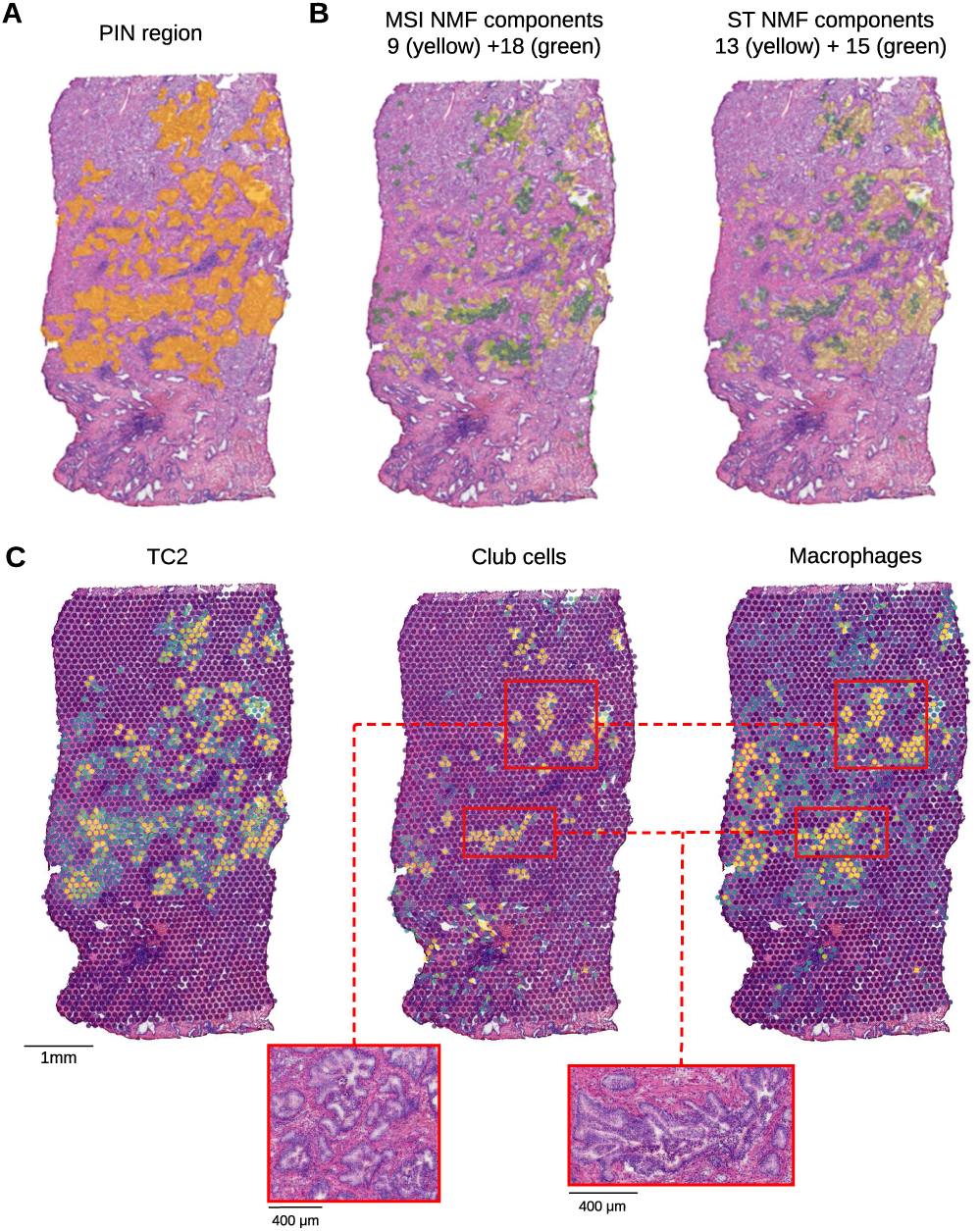
Integration of spatially resolved omics unveils molecular heterogeneity in PIN. **A.** Pathologist annotation of PIN in sample 929_cancer (highlighted in yellow). **B** MSI and ST NMF components correlating with the annotated PIN region in panel A. **C.** Localisation of TC2, club cells and macrophages based on snRNA-seq-deconvoluted ST data. Inserts show H&E stainings of the indicated areas.

### Integrated spatial multi-omics aids in the histological annotation of morphologically ambiguous tissue areas

Integrated SMOx analysis of the entire set of 8 PCa samples overall revealed a strong concordance with the pathologist’s annotations. Nevertheless, in several instances our integrated data-analysis pipeline (based on ST and MSI NMF components) suggested alternative histopathological assignments for areas in the tissues. This was for instance the case in sample 941_normal in which a region initially labelled as PIN exhibited closer molecular concordance with tumour ISUP4 (Fig. 6A**-B**). This observation was further supported by the snRNA-seq data, which indicated the presence of a population of TC1 cells. Furthermore, we observed a match between the top correlated MSI features, predominantly lipids, and genes of the ISUP 4-associated component with the top correlated MSI features and genes of the TC1 cell population (Fig. 6C). Re-examination of this region by the pathologist confirmed the presence of a micro-area of infiltrating prostate adenocarcinoma cells (lower inset box, Fig. 6A**-B**). Another ambiguous tissue area was identified in sample 931_cancer, which, based on the H&E staining of the frozen tissue section was annotated by the pathologist as “other” (Fig. 6D). Based on the ST and MSI NMF components (the latter one showing the strongest spatial correlations), our integrated approach identified this region as a subcluster within the ISUP5 tumour region. snRNA-seq data deconvolution further revealed the presence of two distinct tumour clusters, TC1 highly correlated with ISUP5 and TC2 correlated with the ambiguous region (Fig. 6E**-F**). Histopathological re-evaluation of this region assigned it as cribriform. These results highlight the potential of spatial multi-omics to aid in histopathological annotation of ambiguous tissue areas.

**Figure 6.**
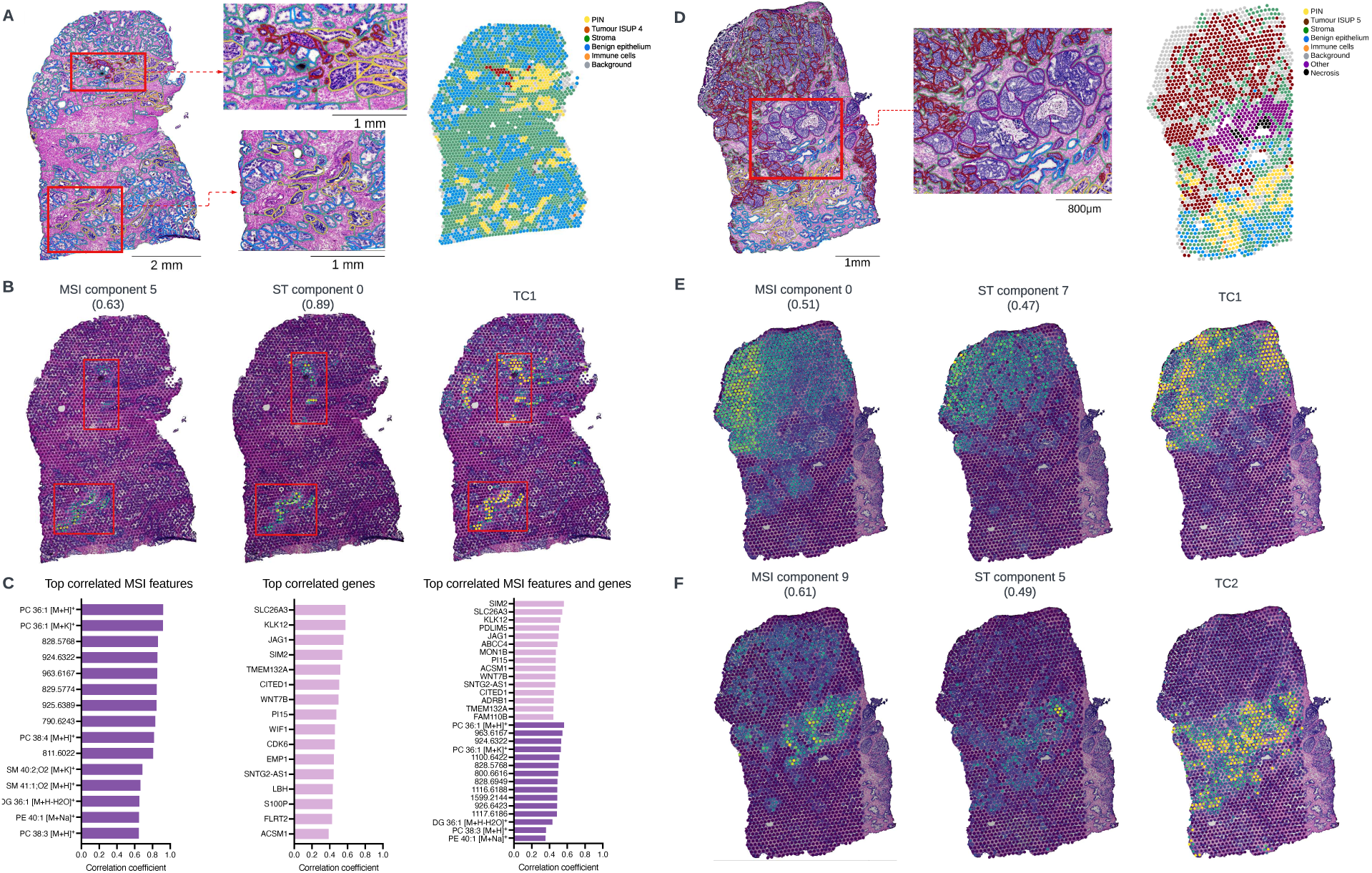
Integrated spatial multi-omics supporting histological annotation of ambiguous tissue areas. **A.** Highly detailed pathologist annotations of a non-malignant prostate sample (941_normal) on the post-MSI H&E-stained tissue section and their spatial distribution projected on the shared coordinate system. Inserts show H&E stainings of tumour ISUP 4 and a morphologically ambiguous region. **B.** Distributions of MSI NMF component 5, ST NMF component 0 and TC1 cell population based on snRNA-seq-deconvoluted ST data. Numerical values (0.63 and 0.89) correspond to the respective correlation coefficient of the MSI and ST NMF with the region annotated by the pathologists as tumour ISUP 4. **C.** Top correlated MSI features and genes of MSI NMF component 5 and ST NMF component 0, respectively. **D.** Highly detailed pathologist annotations of a high-grade prostate cancer sample (931_cancer) on the post-MSI H&E-stained tissue section and their spatial distribution projected on the shared coordinate system. Inserts show H&E stainings of an area annotated by the pathologist as “other”. **E.** Distributions of MSI NMF component 0, ST NMF component 7 and TC1 cell population based on snRNA-seq-deconvoluted ST data. Numerical values (0.51 and 0.47) correspond to the respective correlation coefficient of the MSI and ST NMF with the region annotated by the pathologists as tumour ISUP 5. **F.** Distributions of MSI NMF component 9, ST NMF component 5 and TC2 cell population based on snRNA-seq-deconvoluted ST data. Numerical values (0.61 and 0.49) correspond to the respective correlation coefficient of the MSI and ST NMF with the region annotated by the pathologists as “other”.

### Implementation of the SMOx workflow across the entire prostate cancer sample cohort reveals tumour-specific correlations between transcripts and lipids

Having demonstrated the ability of the integrated SMOx pipeline to extract relevant information from multi-layered molecular expression patterns within a tissue sample, we expanded our analysis to the entire PCa cohort with additional Pearson correlation analysis. Examination of pairs of transcripts and MSI features with correlation coefficients exceeding 0.4 across all 16 biopsies (comprising 8 cancer samples and 8 matching non-malignant samples), allowed us to identify pairs of transcripts and MSI features with a spatial distribution that either correlated or anti-correlated throughout the cohort (Fig. 7A). These included well-established links i.e., between *FASN* expression and PC34:1^44^, confirming the validity of this approach, as well as several novel links illustrating the hypothesis-generating potential of the SMOx pipeline. Next, Cohen’s d values were calculated between tissue types i.e., tumour versus non-tumour tissue (as annotated by pathologists on the H&E microscopy), aiming to detect transcript-MSI feature correlations that significantly changed between these tissue types. One key example is the MSMB -PE 40:2 [M+H]^+^ pair displaying an associated Cohen’s d value of 1.54, indicating a substantial effect size^71^, which in this case implies a considerable difference in spatial co-expression between tumour and non-tumour samples. In fact, as illustrated by a line plot of correlation coefficients for the pair across the 16 biopsies (Fig. 7B) and the actual spatial distributions of the transcript and the lipid in the shared coordinate system of both tumour and non-tumour samples (Fig. 7C), these markers exhibited a clear spatial co-expression in non-tumour samples, which was absent in the tumour samples (quantified with average correlation coefficients of approximately 0.5 and 0, respectively). Specifically, in the non-tumour samples a striking alignment was observed between the gene *MSMB* and lipid PE 40:2 [M+H]^+^, whereas in tumour samples, while *MSMB* continued to exhibit a robust spatial correlation, the lipid PE 40:2 [M+H]^+^ lost its correlation with the benign epithelial cells (highlighted in green). This spatial correlation between PE 40:2 with tumour regions was observed in the majority of tumour samples. While the precise function of this lipid remains elusive, *MSMB* has been previously shown to exhibit a higher expression in normal prostate tissue than in cancerous tissue^72,73^. These observations highlight the hypothesis-generating potential of our SMOx approach.

**Figure 7.**
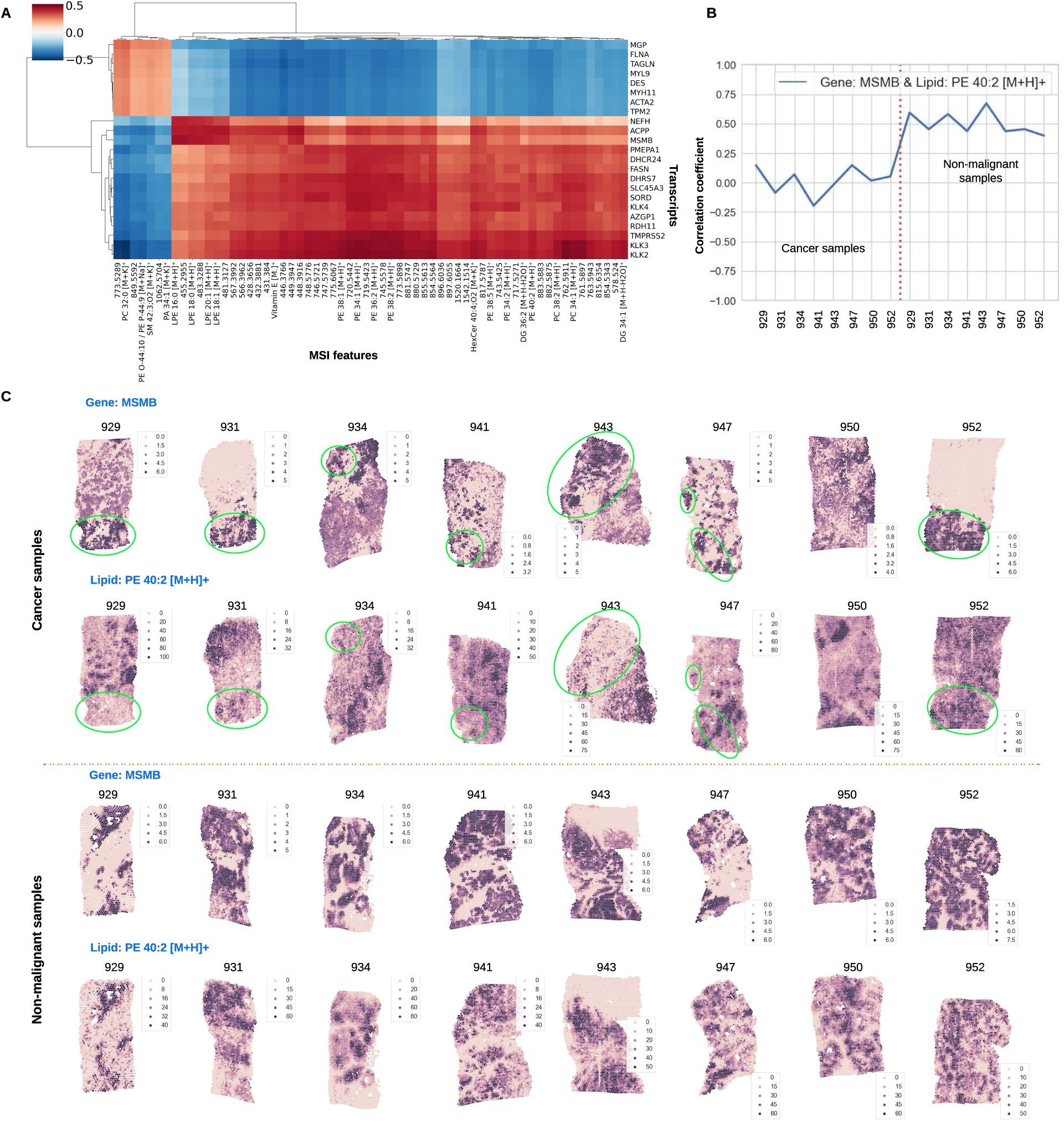
Implementation of the spatial multi-omic workflow across the entire prostate cancer sample cohort. **A**. Heatmap illustrating correlations between genes and MSI features, mostly lipids, where correlation coefficients greater than 0.4 are observed across the entirety of the sample cohort consisting of 16 biopsies (8 cancer samples and 8 matching non-malignant samples). **B.** Line plot depicting the correlation coefficients for the gene/lipid pair MSMB/PE 40:2 [M+H]^+^ across the entirety of the sample cohort. **C.** Spatial distribution of the gene/lipid pair MSMB/PE 40:2 [M+H]^+^ observed across the eight cancer biopsies (upper section) and the corresponding non-malignant samples (lower section). Green polygons indicate regions predominantly composed of benign epithelial cells within the cancer samples.

## Discussion

Spatial multi-omics is progressively establishing itself as a powerful tool to better understand molecular events and its interactions in a native tissue context. One of the key challenges related to this approach is the integration of multiple layers of molecular information that are generally acquired using a range of technologies to optimally capture the molecular information in each layer. Although the ability to conduct diverse spatial approaches, including ST and MSI analyses on a single slide has recently been reported^38^, hitherto, most spatial multi-omics studies are performed on neighbouring tissue sections and generate spatial molecular maps with different spatial resolutions. These factors are further complicated by different tissue deformations due to technology-specific tissue processing, different inter-spot distances and offset of the spots. Our study, which integrates ST and MSI analysis aimed to overcome these hurdles by establishing a data-processing pipeline involving spatial co-registration and granularity matching. Co-registration refers to the mapping of pixels between modalities and is a crucial step in multimodal imaging analysis. Registration of imaging datasets is typically contingent on shared anatomical structures but can be especially challenging when modalities have distinct characteristics, or if measurements are performed at vastly different spatial scales^74^. The former challenge is typically addressed by generating a number of representative images for each modality e.g., by manually selecting from the imaging datasets or by generating images that capture spatial trends in the data by dimensionality reduction methods. The latter, multi-scale challenge refers to the fact that a spatial data point, typically referred to as a pixel or voxel, in one modality can cover a different surface area from that in another modality. These areas can range in the order of magnitude from square nanometres to hundreds of square micrometres. This problem is receiving increasing attention in the spatial biology field, due to the increasing number and complexity of spatial omics applications^75^. A granularity matching step is required to make data points reasonably equivalent between different modalities, typically via an aggregation strategy such as interpolation^8^, averaging, or more advanced methods e.g., Gaussian smoothing^53,76^. In a recent spatial multi-omics study, combining MSI at 15 µm pixel size with multispectral immunofluorescence microscopy^37^, a two-step co-registration was used reporting subcellular accuracy and ROIs from the microscopy projected onto the MSI data, however no granularity matching step was reported. Another study combining MSI and ST using the Visium platform used a spatial resolution of 100 µm, corresponding to the largest modality used, when integrating their data, but do not report whether aggregation was used to account for capture areas not being distributed equally across three modalities^9^. Another study used a weighted nearest neighbour strategy that mechanistically paired Visium spots in the centre of one 50 µm MSI pixel^8^. In contrast to these studies, we used a Gaussian approach that accounts for sample coverage per data acquisition area (Visium spot/MSI pixel). Together with our two-step co-registration tool, a full data analysis pipeline was established that has the ability to deal with the mentioned hurdles in spatial multi-omics data integration.

When applied to a set of clinical PCa samples, downstream integrated analysis revealed a strong spatial concordance amongst the individual modalities as seen in UMAP and NMF analysis. Importantly, agreement between the modalities in this combined approach allows more confidence in defining tissue profiles in spatial atlas approaches, assignment of molecularly heterogeneous regions not discernible by morphology, and potential uses for identifying missed cancerous regions e.g., in a diagnostic context. Using our approach to combine ST, MSI and snRNA-seq analysis on PCa samples, we were able to find lipid and transcript profiles that correlated to specific histological states and to specific cell populations. Based on these correlations between lipids and transcript profiles, we were able to assign unique lipid profiles to specific cell types and disease states and to generate unprecedented insight into the heterogeneity of PCa pathology in the context of a native tissue.

One striking example of the power of our approach was the discovery of a molecularly distinct profile in a PIN region based on both ST and MSI profiling, providing more confidence in the pathological characterisation. Integrated snRNA-seq-based deconvolution revealed the presence of club cells, which recently have been linked to the pathogenesis of PCa^63^. Although the clinical significance of this finding will have to confirmed, our observation of the presence of club cells in a specific pre-malignant histological structure, may point towards that region’s future potential to develop into adenocarcinoma and warrants further investigation.

When applied to the entire cohort of PCa and matched benign samples, we found that spatial transcriptomics and spatial lipidomics data largely followed histomorphological patterns. In addition, combining the two analyses allowed us to identify genetic and lipid expression profiles corresponding to different cell populations and tissue states. Our integrated spatial multi-omics approach was able to identify transcript – MSI feature pairs that correlate or anti-correlate throughout the cohort. Besides well-known associations, including the correlation between *FASN* and PC34:1^44^, which provides validation for our approach, several new correlations were found that warrant future investigation. One striking example was the loss of co-expression of specific lipid-gene pairs such as PE 40:2 [M+H]^+^ and *MSMB* in cancer samples. While the role of *MSMB* as a secreted protein with decreased expression in cancerous tissue is well established^72^, the implications of a change in the abundance of correlating lipid currently remains unknown. Based on transcript – MSI feature pairs and snRNA-seq-based deconvolution of the entire cohort of samples our approach also discovered the presence of tumour cells in a histological region previously assigned as PIN and in a normal tissue section, demonstrating the ability of this approach to aid in pathological annotations. Taken together, these observations illustrate the potential of integrated spatial multi-omics approaches to reveal tissue heterogeneity and to generate new hypotheses as potential starting points for further investigation. Drawing biological conclusions from this concept study should be taken with care in view of the small number of samples that were measured and were selected based on morphological heterogeneity rather than for testing a specific hypothesis. One issue to be further resolved is the presence of batch effects^77^, restricting the comparison of multiple samples to correlation analysis on each sample individually rather than collectively. While various strategies have been proposed to alleviate this issue in both ST and MSI^78^, further method development is required to determine how to mitigate its effects.

As spatial technologies are rapidly evolving and are continuously gaining sensitivity, molecular coverage and spatial resolution, flexible approaches to integrate different layers of spatial and bulk molecular information will become indispensable. Integration approaches, such as the one presented here, will be crucial to better understand the complex interactions between different classes of molecules in a spatial cellular context and to apply this knowledge in diverse fields such as biomarker or target discovery, molecular pathology, and automated tissue annotation.

## Methods

### DATA ACQUISITION

#### Sample collection

Matched prostate tumour and normal samples were obtained from treatment naive high-risk PCa patients (advanced clinical stage and/or biopsy Gleason 8-10 and/or PSA ≥ 20 ng/mL) who underwent radical prostatectomy at University Hospitals Leuven, Belgium. This study was approved by the local Ethics Committee (approval number S54424) and followed the Code of Conduct for Responsible Use of Human Tissue for Research. All patients provided informed, signed consent. After surgery, 6 mm biopsy cores were extracted from the primary tumour site and a matching non-malignant biopsy was taken from the opposite side. The selection of biopsy cores was guided by a urologist and magnetic resonance imaging reports. Immediately after surgery, these biopsy cores were embedded in 3% carboxymethyl cellulose (CMC) (Sigma-Aldrich, cat.no C4888), snap-frozen in isopentane pre-chilled in liquid nitrogen, then stored at −80°C until further processing.

For each prostate biopsy, 10μm thick cryo-sections were obtained at −20°C using a Microm HM525 NX cryostat (Thermo Scientific), followed by H&E staining. Subsequently, two independent expert pathologists performed a thorough examination to verify the presence of tumours in the tumour samples. In this study, a cohort of 8 patients was selected based on detailed pathological information including classification, ISUP grade and morphological features present (see **Supplementary Table 1** for patient-related information). Tissue blocks from the selected patients were subsequently checked for RNA quality. RNA was isolated from ten 10µm sections from the selected tissue blocks using Direct-zol RNA Miniprep (Zymo Research, cat.no. R2050), and RIN score was determined using a Bioanalyzer RNA 6000 Nano kit (Agilent, cat.no 5067-1511) in combination with the Agilent Bioanalyzer software. All samples had a RIN > 7. 10μm thick sections were collected on ITO-coated glass slides (Delta Technologies, Loveland, USA) to enable matrix-assisted laser desorption/ionisation (MALDI) MSI analysis, and adjacent tissue sections of 10μm was mounted onto pre-equilibrated 10x Genomics Visium Gene Expression slides (10x Genomics, Pleasanton, CA, USA) for ST analysis. Finally, three to five tissue sections with a thickness of 100μm were collected in a pre-cooled centrifuge tube for subsequent single-nuclei RNA sequencing (snRNA-seq) analysis.

#### Spatial transcriptomics sample preparation and data acquisition

Tissue preparation for Visium ST analysis was performed according to the Tissue Preparation Guide (CG000240 rev C, 10x Genomics), except for embedding in 3% CMC which was conducted to stabilise the tissue for both ST and MSI analysis. Methanol fixation, H&E staining and imaging were performed according to the recommended protocol (CG000160 rev B, 10x Genomics), and subsequent cDNA and library preparation was performed according to the Visium Spatial Gene Expression Reagent Kit User guide (CG000239 rev D, 10x Genomics). The optimal tissue permeabilization time was determined upfront to be 20 minutes, following the Visium Spatial Gene Expression Reagent Kit -Tissue Optimization User guide (CG000238 rev A, 10x Genomics). Final libraries were sequenced to at least 50.000 reads/covered spot using a NextSeq2000 platform (Illumina, San Diego, CA, US).

#### Mass spectrometry imaging sample preparation and data acquisitions

Tissue sections were coated in 2,5-dihydroxybenzoic acid (DHB, AK Scientific, CA, USA) matrix using a sprayer robot (TM-Sprayer, HTX Technologies, LLC, Chapel Hill, NC, USA). DHB was dissolved in a 1:1 mixture of methanol:chloroform (v:v) at a concentration of 20 mg/mL. 12 layers of matrix solution were deposited using a flow rate of 0.12 mL/min, 10 psi of N2 pressure, nozzle temperature of 30 °C, and a 1200 mm/min velocity.

All data was acquired using an Orbitrap Elite mass spectrometer (Thermo Fisher Scientific GmbH, Bremen, Germany) coupled to an intermediate pressure MALDI source (Spectroglyph LLC, WA, USA) as described previously^79^. A frequency tripled Nd:YLF laser (Explorer One, Spectra Physics, Mountain View, CA) emitting at 349 nm operating at 300 Hz was used for MALDI. The laser was operated using a diode current of 1.80 A and pulse energy fine-tuned using an external attenuator (PowerXP, Altechna, Vilnius, Lithuania) positioned immediately in front of the laser to yield a post-attenuation pulse energy of 1.2 µJ. A frequency quadrupled laser Nd:YAG laser emitting at 266 nm and operating at 300 Hz (NanoDPSS, Litron Lasers, Warwickshire, England) was used for laser post-ionisation (MALDI-2). This setup has been described in detail previously^80^. MALDI-2 laser pulse energy was adjusted using the internal attenuator to be 600 µJ just prior to entering the ion source. Laser pulse energy measured using a calibrated energy sensor (QE12HR-H-MB-D0, Gentec-EO, Quebec, Canada). Both lasers were externally triggered using a pulse/delay generator (QC9200, Quantum Composers, Bozeman, Montana) such that the time delay between the MALDI and MALDI-2 laser pulse was 20 µs. The mass spectrometer was operated in positive ion mode using an ion injection time of 250 ms, automatic gain control turned off, m/z range 350-2000 and a nominal mass resolution of 120,000 @ m/z 400. The pixel size was 30 µm.

All raw files were first recalibrated with Thermo Scientific Xcalibur RecalOffline using ion signals of [Vitamin E]^+^ at *m/z* 430.38053, [PC(34:1)+K]^+^ at *m/z* 798.54096 and a background PDMS peak at *m/z* 371.10124. Data was then converted to imzML format using Image Insight software (Spectroglyph LLC, Kennewick, WA, USA).

#### Tissue Staining and Histopathology Annotations

Tissue staining using H&E was conducted on tissue sections following MSI analysis. The matrix was first removed from the slide by rinsing in methanol. For the H&E staining, the glass slides were immersed in 95% ethanol, 70% ethanol and H2O for 2 minutes each and then immersed in haematoxylin for 3 minutes. Slides were then washed with running tap water for 3 minutes and immersed in eosin for 30 seconds following another washing step with water for 3 minutes and immersion in 100% ethanol for 1 minute and then in xylene for 30 seconds. The tissues were air-dried and mounted with coverslips. Optical images were acquired using a ZEISS Axio Scan.Z1 Slide Scanner (Carl Zeiss AG, Jena, Germany) with 40x objective, pixel size 0.22 µm* 0.22 µm. The files were exported as .TIFF images.

The digitalized H&E images were annotated and verified by prostate cancer specialists from the Department of Pathology, University Hospitals Leuven and Weill Cornell Medical College, in a blinded fashion using the web-based annotation tool ‘Annotation studio’ (Aspect Analytics NV, Genk, Belgium). Pathologists annotated the whole slides in detail by placing 14 different labels ‘Inflammatory infiltrate’, ‘Normal epithelium’, ‘Stroma’, ‘Necrosis’, ‘Atrophy’, ‘Cribriform’, ‘PIN’, ‘Nerve’, ‘Tumour ISUP 5’, ‘Tumour ISUP 4’, ‘Tumour ISUP 3’, ‘Tumour ISUP 2’, ‘Tumour ISUP 1’, and ‘Other’.

#### Single cell RNA sequencing sample preparation and data acquisition

Tissue sections (2-3×100 µm) adjacent to those used for spatial transcriptomics and lipidomics sections were collected in a microcentrifuge tube and subjected to single nuclei dissociation. For this, 0.3mL Tween with Salts and Tris (TST) buffer (96mM NaCl (Sigma-Aldrich, cat.no. 59222C), 10mM Tris-HCl pH 7.5 (ThermoFisher, cat.no. 15567027), 1mM CaCl2 (Merck, cat.no. 21115), 21mM MgCl2 (Sigma-Aldrich, cat.no M1028), 0.03% Tween-20 (Biorad, cat.no 1662404), 0.01% BSA (New England Biolabs, cat.no B9000S) and 0.2U/µl RNAsin (Promega, cat.no N2615)) were added to the sections and the tissue was cut into small pieces using scissors. The tissue was subsequently transferred to a KIMBLE Dounce tissue grinder (Sigma-Aldrich, cat.no D8938), together with 0.7mL TST that was used to wash the original tube. The tissue was homogenised by 10 strokes of the large clearance pestle and 10 strokes of the small clearance pestle. The homogenate was subsequently filtered using a 40µm cell strainer (pluriSelect, cat.no. 43-50040-51) and the tissue grinder and filter were washed with Salts and Tris (ST) buffer (96mM NaCl (Sigma-Aldrich, cat.no 59222C), 10mM Tris-HCl pH 7.5 (ThermoFisher, cat.no 15567027), 1mM CaCl2 (Merck, cat.no. 21115), 21mM MgCl2 (Sigma-Aldrich, cat.no M1028) and 0.2U/µl RNAsin (Promega, cat.no. N2615)). Filtered nuclei were pelleted twice for 5 minutes at 500g at 4°C, first followed by resuspension in 500 µl ST buffer, and second followed by resuspension in 50µl of Chromium loading (CL) buffer (PBS + 0.04% BSA + 1U/µl Sigma Protector RNase Inhibitor (Millipore Sigma, cat.no. 3335399001). Nuclei were counted using a PhotonSlide™ (Logos Biosystems, South Korea) with Acridine Orange & Propidium Iodide Cell Viability assay (Westburg, cat.no. LB F23001) on a LUNA-FL™ Dual Fluorescence Cell Counter (Logos Biosystems). Nuclei were counted as the ‘dead’ cell fraction, and the suspension was diluted to ∼1.000.000 nuclei/mL in CL buffer.

Single nuclei suspensions were used for single-nuclei gene expression analysis using the Chromium Next GEM Single Cell 3’ Reagent Kits v3.1, according to the user guide (CG000315 Rev C). 7000-8000 nuclei were loaded per sample, and final libraries were sequenced to at least 25.000 reads/cell using a NextSeq2000 platform (Illumina, San Diego, CA, US).

### DATA PREPROCESSING

#### Spatial transcriptomics data pre-processing

FASTQ-files were mapped to the GRCh38 human reference genome and spatially projected using Space Ranger v1.1.0 (10x Genomics). Before merging the datasets, ambient RNA was removed using an adjusted version of SpotClean v1.1.1 where all relevant genes are maintained^81^. Next Scanpy v.1.9.1 was used for quality control and processing^82^. Spots with less than 700 genes and 1000 reads were excluded, while genes were filtered for expression in more than 100 spots. The Scanpy toolkit was used to perform downstream processing per sample, including normalisation with the Scran R package (v1.26.1), log transformation and variable gene detection. The number of significant principal components for Leiden clustering and UMAP dimensionality reduction was determined using bootstrapping with a maximum of 30 components. Clustering resolutions ranging from 0.1-1.0 were assessed for stability with steps of 0.1 using clustree v0.4.2^83^.

#### Spatial lipidomics data preprocessing

The mass spectrometry imaging data was preprocessed by performing TIC normalization, peak picking and subsequent rebinning. Peak picking was performed on the mean spectrum across all datasets. Peaks in the mean spectrum with an intensity below 0.005% of the base peak were discarded, resulting in a total of 12510 retained peaks. Intensities were binned per spectrum using a 5 ppm window around the selected peaks.

#### Spatial multi-omics integration pipeline

As the ST and MSI data were acquired from two serial sections, the spatial lipidomics and transcriptomics data were indirectly registered to each other by their respective H&E microscopy images. This strategy of co-registering multiple sections via one proxy section (i.e., microscopy) is inevitable in applications where multiple sections are involved. Before co-registration, representative images from the MSI data were generated for guiding the later registration step. These representative images were created via applying dimensionality reduction methods, such as NMF and UMAP. The MSI data was co-registered with its respective H&E microscopy images via a non-rigid registration workflow^84^ (Aspect Analytics NV). Next, the MSI-slide H&E was co-registered to the H&E from the ST Visium slide. No additional co-registration is required to link the ST data and ST-slide microscopy as their spatial reference is inherent to the Visium ST data format. Finally, the MSI was directly linked with the ST data via a shared spatial coordinate system. MSI data was acquired at a spatial resolution (pixel size) of 30 µm, whereas the ST data was obtained with spot diameters of 55 µm. Multiple MSI pixels were aggregated into a single representative spectrum to match the ST spot via a Gaussian weighting algorithm^53^.

#### Morphological feature extraction from microscopy data

Morphological features from high resolution H&E microscopy image of the MSI section were extracted using a pre-trained model that was adapted originally from the SimCLR model introduced by Chen et al^85^. The model tries to maximize the agreement between two stochastically augmented views of the same image via a contrastive loss function. Ciga et al. later applied this SimCLR model with minor modifications and pre-trained it on 57 multi-organ histopathology datasets without any labels^86^. These 57 histopathology datasets contained microscopy images with various types of staining and resolutions, which helped the model learn features with better quality. We applied this pre-trained model on high-resolution H&E-stained microscopy data, acquired at a 40x magnification, and extracted the morphological features using overlapping windows of 512 × 512-pixel patches of the image centred on each spot. Thus, each patch covers approximately 128×128 µm^2^, which is similar in size to the patches extracted from histopathology datasets that the model was pre-trained on by Ciga et al^86^.

#### Dimensionality reduction via non-negative matrix factorization

We applied NMF to reduce the high dimensionality of our MSI and ST data. NMF has been widely used in both fields^87–90^ as it has the non-negativity constraint and can generate interpretable parts-based representation. We used NMF to get a more in-depth understanding of each modality in a fully unsupervised manner, based on their resulting distinct spatial distributions and associated spectral or gene expressions. In short, NMF factorizes the matrix X into two non-negative matrices X=WH, where W represents the loading matrix and H is the score matrix which represents the data in lower dimensions. For example, in the case of MSI data, the lipid spectral signature of each pixel can be approximated by the additive linear combination of the column vector (spectral basis element) from W, weighted by H, which describes the contribution of each basis element to the spectrum at each pixel. We implemented NMF by the Python package sklearn.decomposition.NMF. As both MSI and ST data are known to follow Poisson distribution, we used Kullback–Leibler divergence (KL-NMF) as the cost function to incorporate Poisson noise. Multiplicative Update was used as the solver. The number of components was selected as 20.

#### Correlation analysis

We applied Pearson correlation by Python package numpy.corrcoef and scipy.stats.pearsonr to measure the relationship between two datasets. The output correlation coefficient ranges from -1 to 1 with 0 implying no correlation. Correlations of -1 or 1 represent an exact linear relationship (negative or positive).

#### Cohen’s d

We calculated the Cohen’s d value for gene-lipid correlations across different tissue types. Cohen’s d is defined as the difference between two means divided by the pooled standard deviation for the data. The resulting Cohen’s d values were ranked to find the most different gene-lipid correlations between normal and tumour tissue types. Using this approach, a set of changes in correlation were identified. We firstly calculated the Cohen’s d value for gene-lipid correlations across different tissue types via computing their mean correlations in 8 tumour samples and 8 ‘normal’ samples separately.

#### Processing of the single-nuclei RNA-sequencing data and cell type deconvolution

FASTQ-files were mapped to the GRCh38 human reference genome with Cell Ranger v6.1.2 (10x Genomics). Before merging the datasets, ambient RNA was removed using SoupX^91^, while scDblFinder^92^ was used to indicate doubles. Next Scanpy v.1.9.1 was used for quality control and processing^82^. Only cells with the following criteria were considered for further analysis: more than 500 uniquely expressed genes, less than 10% of the UMI counts mapping to mitochondrial sequences and less than 5% of the UMI counts assigned to ribosomal genes.

All genes that were not expressed in at least 100 cells were not considered in the downstream analysis. The Scanpy toolkit was used to perform downstream processing per sample, including normalisation, log transformation and variable gene detection. The number of significant principal components for Leiden clustering and UMAP dimensionality reduction was determined using bootstrapping with a maximum of 60 components. Clustering resolutions ranging from 0.1-1.0 were assessed for stability with steps of 0.1 using clustree v0.4.2^83^. Cell types and states were assigned to the clusters using marker genes obtained from the PanglaoDB^93^ database (version of 27/03/2020), the canonical markers from Song *et al.*^63^ and the NMF signatures obtained from the analysis of matching ST samples. Copy-number landscapes were inferred from the transcriptomic data with inferCNV^94^ v.1.3.3 to validate clusters annotated as tumorigenic. Integration of the samples was performed in a semi-supervised manner using scANVI^95^. First the transcriptome data was integrated using scVI^96^ (v.0.14.5, n_hidden=128, n_latent=50, n_layers=2, dispersion=’gene-batch’) with correcting batch effects and removing unwanted source of variations (total counts, percent mitochondrial genes, and percent ribosomal genes for ‘continuous_covariate_keys’). After training the model for 400 epoch, scANVI using the cluster labels obtained at single-cell level, was trained for an additional 10 epochs. The integrated latent embedding generated by scANVI was used for downstream analysis (clustering and visualisation). For gene-level analyses, uncorrected counts were used. Spot deconvolution was performed using the integrated single-nuclei data per patient and Cell2Location^46^. The lipid metabolism gene sets of the KEGG pathway database^97^ and the ‘score genes’ function of scanpy were used to asses pathway activities.

#### Lipid Annotation/Identification

Lipids were annotated with mass accuracy of 3 ppm based on the LipidMaps guidelines on shorthand notation for lipid structures published by Liebisch et al.^98,99^ For the glycerophospholipids (GPL), the following shorthand notations of lipid classes were used: PC – phosphatidylcholines, LPC – lysophosphatidylcholines, PC O – phosphatidylcholine ethers, PC P – phosphatidylcholine plasmalogens, PE – phosphatidylethanolamines, LPE – lysophosphatidylethanolamines, PE O – phosphatidylethanolamine ethers, PE P – phosphatidylethanolamine plasmalogens, PS – phosphatidylserines, PI – phosphatidylinositols. For the sphingolipids (SPL), the following class name abbreviations were used: Cer – ceramides, SM – sphingomyelins, and Hex(n)Cer – hexosyl ceramides, where n refers to the number of hexosyl units. Cholesterol is referred to as ‘Chol’, cholesteryl esters as CE, acylcarnitines as CAR, monoacylglycerols as MG, and diacylglycerols as DG. Lipids are annotated at the lipid species level, namely the lipid class abbreviation is followed by a total number of carbons in fatty acyl(s) rest(s), colon, and total number of double bonds in fatty acyl rest(s). For example, PC 36:3 refers to phosphatidylcholine with a total number of carbons in both fatty acyls equal to 36 and total number of double bonds equal to 3. The same principle was used for annotation of signals corresponding to sphingolipids, but additionally the total number of oxygen was indicated after semi-colon, for example, SM 36:2;O2 refers to a sphingomyelin with sum composition of carbon atoms in N-linked fatty acyl rest and sphingoid base of 36, the number of double bonds equal to 2, and total number of O-atoms equal to 2. CE 16:0 means cholesteryl ester with palmitic acid attached to cholesterol via an ester bond. In MALDI-2 mode cholesteryl esters form dimers as showed recently by Bowman et al.^65^ For m/z values annotated as dimers of cholesteryl esters, the following shorthand notations were used: 2CE for cholesteryl ester dimer, followed by total number of carbons in both fatty acyls, colon, and total number of double bonds in both fatty acyls e.g., 2CE 38:4.

## ACKNOWLEDGEMENT

We thank all patients that participated in this study.

WZ was supported by a Baekeland PhD grant from Flanders Innovation and Entrepreneurship (VLAIO) - HBC.20192204. XS is a recipient of a PhD fellowship (SB/1S49218N) from the Research Foundation – Flanders (FWO) and was in part supported by Opening the Future (OtF). SRE acknowledges support from the Australian Research Council (FT190100082). TS acknowledges support through the Australian Government Research Training Program Scholarship. SV and VdL were sponsored by an FWO PhD Followship strategic basic research (1S93320N and 1157918N, respectively). KV, SK and TV are supported by KU Leuven (IDN/19/039, Opening the Future (OtF)). This work was supported by a grant from the Belgian Foundation Against Cancer (STK), a Focus Group grant from the LKI Fund for Innovative Cancer Research (FIKO), the KU Leuven Opening the Future campaign, KU Leuven Incubation financing Lipometrix, a grant PRISMO from the Flemish Resilience Plan, FWO-EOS projects 30837538 (DECODE), G0F6718N (SeLMA), FWO-SBO projects S001623N (LIPOMACS) and S005319N, KU Leuven Research Fund projects C14/21/095, BOF/23/064, C16/15/059, C3/19/053, C24/18/022, C3/20/117, C3I-21-00316), US-DOD project W81XWH-19-1-0566, Industrial Research Fund (Fellowships 13-0260, IOFm/16/004) and several Leuven Research and Development bilateral industrial projects, Infrastructure projects TOP/23/014, IO12220N and I013218N, TBM Project T001919N; PhD Grant (SB/1SA1319N), the Flanders AI Research Program, VLAIO CSBO (HBC.2021.0076), Baekeland PhD (HBC.20192204), European Research Council under the European Union’s Horizon 2020 research and innovation programme (ERC Adv. Grant grant agreement No 885682) and EU Interreg EMR23 EURLIPIDS, Stand up to Cancer / Kom op tegen Kanker - the Flemish cancer society, CM (Christelijke Mutualiteit), and private donations. Figure 1 was created with <u>BioRender.com</u>.

## Notes

### Competing Interest Statement

Competing Financial Interests:
A.L., and M.J.Q.M. are employed by Aspect Analytics NV.
M.C. and N.V. are shareholders of Aspect Analytics NV.
T.V. is co-inventor on licensed patent WO/2014/053664 (High-throughput genotyping by sequencing low amounts of genetic material).
The remaining authors declare no competing interests.

