## Supplementary Information for "Integration of Multiple Spatial Omics Modalities Reveals Unique Insights into Molecular Heterogeneity of Prostate Cancer"

### Supplementary Table 1

**Clinical data for the eight patients used in the SMOx study**

| | Patient ID | 929 | 931 | 934 | 941 | 943 | 947 | 950 | 952 |
|---|---|---|---|---|---|---|---|---|---|
| | Age at surgery | 73 | 61 | 72 | 64 | 72 | 62 | 72 | 69 |
| Patient characteristics | BMI | 26,21 | 25,14 | 27,77 | 27,7 | 28,37 | 21,13 | 36,56 | 30,55 |
| | date surgery | 07/09/2020 | 17/09/2020 | 18/11/2020 | 08/02/2021 | 11/02/2021 | 04/03/2021 | 11/03/2021 | 13/04/2021 |
| | Medication with an influence on lipids | 1 | 0 | 1 | 1 | 1 | 0 | 0 | 0 |
| | Which? | Crestor 10mg | NA | Crestor 20mg | Crestor 20mg + Ezetimibe 10mg | Atorvastatine 10mg | NA | NA | NA |
| | iPSA | 8,7 | 16,2 | 19 | 14,6 | 4,81 | 8,04 | 24 | 3,79 |
| | cT-stage | 1 | 2b | 1 | 3a | 1 | 1 | 3a | 2a |
| Pre-OP characteristics | Biopsy GS | 4+4=8 | 4+4=8 | 4+4=8 | 4+3=7 | 3+4=7 | 3+5=8 | 4+5=9 | 3+4=7 |
| | Biopsy ISUP stage | 4 | 4 | 4 | 3 | 2 | 4 | 5 | 2 |
| | Neoadjvuant therapy | 0 | 0 | 0 | 0 | 0 | 0 | 0 | 0 |
| | pT-stage | 2c | 3b | 3a | 3b | 2c | 2c | 3a | 3a |
| | Lymph node invasion | 0 | 1 | 0 | 1 | NA | 0 | 0 | 0 |
| | Lymphovascular invasion | 0 | 1 | 1 | 0 | 0 | 1 | 0 | 0 |
| Pathology of specimen | Perineural invasion | 1 | 1 | 1 | 1 | 1 | 1 | 1 | 1 |
| | Section margins | 0 | 0 | 0 | 1 | 0 | 0 | 0 | 0 |
| | pGS | 3+4=7 | 4+5=9 | 4+3=7 | 4+5=9 | 4+3=7 | 3+4(+5)=7 | 4+3(+5)=7 | 4+3=7 |
| | pISUP | 2 | 5 | 3 | 5 | 3 | 2 | 3 | 3 |
| | Undetectable post-OP PSA | 1 | 0 | 0 | 0 | 1 | 0 | 0 | 1 |
| | Adj RT | 0 | 1 | 1 | 1 | 0 | 1 | 0 | 0 |
| | RT because of BCR or rising PSA | NA | 0 | 1 | 0 | NA | 1 | NA | NA |
| | Time from surgery till RT | NA | 4,8 | 13,43 | 3,33 | NA | 26,73 | NA | NA |
| | PSA nadir | NA | 0 | 0 | 0 | NA | NA | NA | NA |
| | Adj ADT | 0 | 1 | 1 | 1 | 0 | 1 | 0 | 0 |
| | Time from surgery till ADT | NA | 4,8 | 13,43 | 3,33 | NA | 26,73 | NA | NA |
| Follow-up | BCR (PSA ≥ 0,2) | 0 | 0 | 0 | 0 | 0 | 1 | 0 | 0 |
| | Time from surgery till BCR | NA | NA | NA | NA | NA | 24,7 | NA | NA |
| | Clinical recurrence? | 0 | 0 | 0 | 0 | 0 | 1 | 0 | 0 |
| | Time from surgery till CR | NA | NA | NA | NA | NA | 24 | NA | NA |
| | Last PSA | 0 | 0,15 | 0 | 0 | 0 | 0,23 | 0,1 | 0 |
| | Date last PSA | 26/04/2023 | 25/04/2023 | 31/05/2023 | 09/01/2023 | 13/03/2023 | 24/03/2023 | 04/04/2023 | 22/03/2023 |
| | Follow-up (months) | 31,67 | 31,3 | 30,43 | 23,07 | 25,1 | 26,73 | 24,8 | 23,33 |
| | Alive with no evidence of disease | 1 | 0 | 1 | 1 | 1 | 0 | 0 | 1 |
| | Remarks? | / | No T level | Testosterone 0,133 nmol/L | Testosterone 12 ng/dL | / | RT still going | PSA rising | / |

Supplementary Figure 1

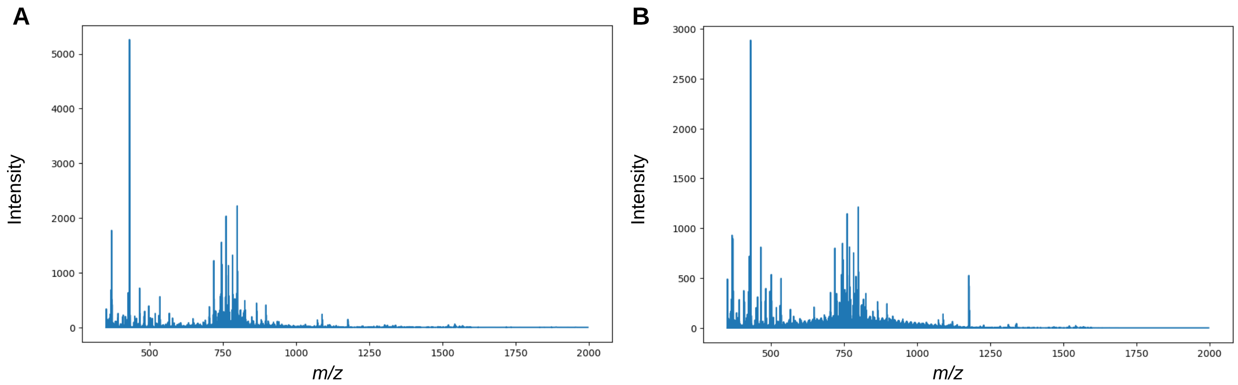


**Supplementary Figure 1. Averaged MALDI-2 spectra. A.** Averaged MALDI-2 spectrum of all cancer samples. **B.** Averaged MALDI-2 spectrum of all matching non-malignant samples.

Supplementary Figure 2

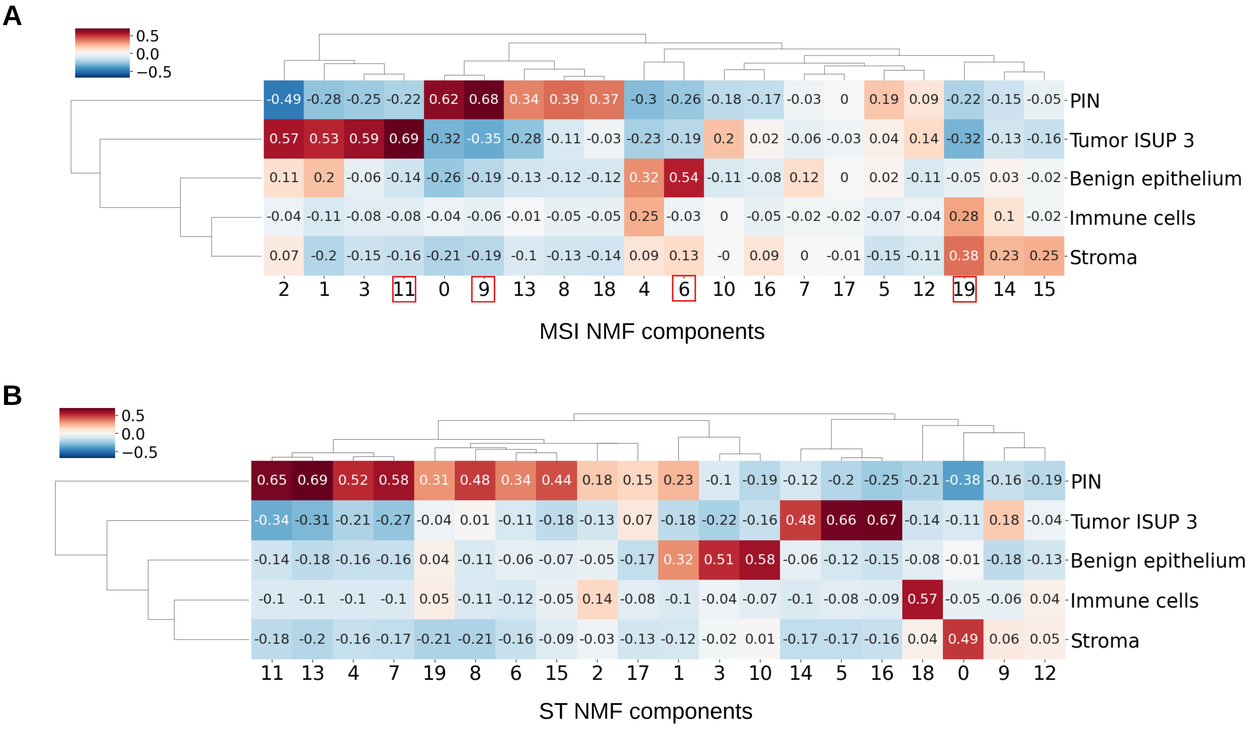


**Supplementary Figure 2. Correlation heatmaps comparing pathology annotations versus spatial omics expression data. A.** Correlation between MSI NMF components and the five histologically distinct regions from sample 929_cancer. **B.** Correlation between ST NMF components and the five histologically distinct regions from sample 929_cancer.
